# Testosterone improves muscle function of the extensor digitorum longus in rats with sepsis

**DOI:** 10.1101/850636

**Authors:** Jinlong Wang, Tong Wu

## Abstract

Among patients with Intensive care unit-acquired weakness (ICUAW), skeletal muscle strength often decreases significantly. This study aimed to explore the effects of testosterone propionate on skeletal muscle using rat model of sepsis. Male SD rats were randomly divided into experimental group, model control group, sham operation group and blank control group. Rats in experimental group were given testosterone propionate 2 times a week, 10 mg/kg for 3 weeks. Maximal contraction force, fatigue index and cross-sectional area of the extensor digitorum longus (EDL) were measured. Myosin, IGF-1, p-AKT and p-mTOR levels in EDL were detected by Western blot. Histological changes of the testis and prostate were detected by hematoxylin and eosin staining. We found that maximal contraction force and fatigue index of EDL in experimental group were significantly higher than in model control group. Cross sectional area of fast MHC muscle fiber of EDL in group was significantly higher than in model control group. The levels of myosin, IGF-1, p-AKT and p-mTOR of EDL in experimental group were significantly higher than in model control group. In addition, no testicle atrophy and prostate hyperplasia was detected in experimental group. In conclusion, these results suggest that testosterone propionate can significantly improve skeletal muscle strength, endurance and volume of septic rats, and the mechanism may be related to the activation of IGF-1/AKT pathway. Moreover, testosterone propionate with short duration does not cause testicular atrophy and prostate hyperplasia in septic rats. Therefore, testosterone propionate is a potential treatment for muscle malfunction in ICUAW patients.

## Introduction

Intensive care unit-acquired weakness (ICUAW) is common in critically ill patients who have decreased muscle strength and weakened peripheral nerve excitability.^1^ At present, it is generally believed that the occurrence of ICUAW is related to multiple factors, including sepsis, multi-organ failure and mechanical ventilation, among which sepsis is the most prominent.^2^ ICUAW affects approximately 40% of critically ill patients, and even 50-100% patients with severe sepsis.^3^ Therefore, prevention and treatment of ICUAW has become an urgent medical issue.^4^

Testosterone is a major male hormone applied in clinical practice in patients with severe burns and other weaknesses to improve muscle function and quality of life.^5^ Testosterone treatment leads to a significant dose-dependent increase in muscle function, myonuclei and satellite cells aggregation in the muscles of the elderly. ^6^ In animal experiments, testosterone improved the muscle function of paraplegia, limb fracture, and castration.^6,7^ Therefore, determining whether the appropriate testosterone supplements can be given to ICUAW patients is interesting and it needs further studies.

ICUAW patients often show skeletal muscle atrophy. One major role of testosterone is to promote the synthesis of skeletal muscle protein by acting on a series of cellular signals through androgen receptors.^9,10^ In addition, testosterone promotes the proliferation of vascular endothelial cells to increase local blood and oxygen supply of muscles, ultimately increasing muscle volume, strength and endurance.^11^ Clinical and animal studies have shown that testosterone can significantly increase the expression of tissue and circulating insulin-like growth factor 1 (IGF1), which activates Akt/m-TOR pathway to regulate skeletal muscle.^12,13^

Therefore, this study aimed to investigate the effect and possible mechanism of testosterone on muscle weakness in rat model with sepsis.

## Materials and methods

### Animal

Animals experiments were conducted at Chongqing Medical University in accordance with “Chongqing Administrative Measures on Experimental Animals” and approved by the ethics committee of Chongqing Medical University. Healthy male SD rats were purchased from the Animal Experiment Center of Chongqing Medical University (license number scxk-2017-0001) and raised in an SPF level animal house. Rats were fed standard feed at room temperature (20 °C), with a circadian rhythm of twelve hours of day alternated with twelve hours of night.

The rats were randomly divided into four groups (n=8): experimental group, model control group, sham operation group, and blank control group. The rats were anesthetized with pentobarbital salt solution (40 mg/kg) via intraperitoneal injection, and then cecal ligation and perforation was performed in experimental group and model control group to establish the animal model of sepsis. Anesthesia and abdominal operation were performed without ligation and perforation in sham operation group. The blank control group did not receive any operation. The 8th day after surgery, hind limb was fixed, and a 1 ml sterile empty needle was used for vertical puncture into the muscle. Sterile cotton balls were used for compression and hemostasis after injection, and the hind limbs on both sides were injected with testosterone propionate (Guangzhou Baiyunshan pharmaceutical) twice a week, 10 mg/kg for 3 weeks for rats in experimental group; rats in model control group were injected with same amount of soybean oil with the same injection method and date. Rats in sham group and blank control group were not injected.

### Determination of serum testosterone

One ml blood was collected from the rats by the orbital blood collection method. The serum testosterone concentration was determined by radioimmunoassay using ^125^I testosterone radioimmunoassay kit for rat (Beijing Furuirunze Biotechnology Co, LTD).

### Maximum contractile force, contraction time, relaxation time and fatigue index

Rats were anesthetized with pentobarbital and were placed on a mat heated to 37 °C. The sciatic nerve was dissected intact through a left hip incision, and small branches of the sciatic nerve innervating the hip muscle were cut off, leaving only the anterior tibial nerve. A longitudinal incision was made in the middle of the foot, and the extensor digitalis longus (EDL) was disassociated along the tendon, while all blood vessels were preserved. The rats were placed in prone position, with the foot fixed at 90° of the ankle, and the femur fixed with clamps at an angle of approximately 100° between the femur and the tibia.^14^ The tendon of EDL with the tension transducer was connected with an inelastic thread, and the sciatic nerve was gently placed on the hook of a bipolar electrode. A macro-adjuster was used to extend the length of EDL gradually by 1 mm each time until the contraction force decreased. The length of EDL corresponding to the maximum contraction force was the optimal initial length. EDL was relaxed and returned to the state of natural relaxation, the optimal initial length was recorded, and all measurements were determined under this length. The contraction time is defined as the time from the beginning of the contraction to the maximum peak, and the relaxation time is defined as the time from the peak to the baseline. The percentage of the final contraction force to beginning contraction force of EDL was named the fatigue index of the rat toe extensor.^15^

### Muscle, testicle and prostate specimen collection

Rats were sacrificed by high-dose pentobarbital (100 mg/rat) and high carbon dioxide. The standard of death was loss of consciousness, respiratory arrest and electrocardiogram. EDL of the right leg was cut off and frozen at −80 °C, while the middle segment of the left EDL, testicle and prostate tissue were cut off and fixed in 4% paraformaldehyde.

### Western blot analysis

EDL was homogenized in ice-cold lysis buffer (pH 7.4) containing 50 mM Tris-base, 1% (v/v) Triton X-100, 0.25% (v/v) sodium desoxycholate, 150 mM NaCl, 1 mM EDTA, 1 ug/mL leupeptin, 1 ug/mL aprotinin, 5 mM DTT, and 1 mM PMSF. All homogenates were centrifuged at 12,000 x g, 4 °C for 10 min, and the supernatants were centrifuged at 100,000 x g at 4 °C for 1 h. The protein concentration was determined using the Bradford Protein Assay Kit (Beyotime, China). Soluble proteins (20 μg) were used for Western blotting with primary antibodies against AKT1/2/3 (rabbit monoclonal antibody; EPR16798 Abcam; 1:10,000), mTOR (rabbit monoclonal antibody; Y391 Abcam; 1:2,000), mTOR phosphor S2481 (rabbit monoclonal antibody; EPR427(N) Abcam; 1:5,000), AKT1/2/3 phospho Y315 + Y316 + Y312 (rabbit polyclonal antibody; Abcam; 1:1,000) and fast myosin skeletal heavy chain (rabbit polyclonal antibody; Abcam; 1 μg/ml). and goat anti rabbit IgG (Abcam, 1:5,000). Chemiluminescence was performed using a super ECL plus (Beyotime, China), and the blots were exposed to a Gel Imaging System (Bio-Rad, America).

### Histological analysis

The frozen EDL tissues were sliced with a Cryostat Microtome (Leica, Germany). Immunohistochemical analysis of the sections was performed using fast myosin skeletal heavy chain antibody (Abcam, ab91506, UK), and the staining was analyzed with Image Pro Plus 6 software. The testicle and prostate tissues were sectioned for hematoxylin and eosin (HE) staining, and observed under an optical microscope (Olympus Optical, Japan).

### Statistical analysis

All data was expressed as Mean ± standard deviation (SD) and analyzed by SPSS 21.0 software. For data with normal distribution, one-way ANOVA was used to compare the difference, and for data not in normal distribution, Mann-Whitney U test was used to compare the difference. P < 0.05 was considered significant.

## Results

### Changes in body weight and serum testosterone concentration in each group

Preoperative body weight of rats among each group showed no significant difference. Body weight of rats in experimental group and model control group significantly decreased compared to sham group and blank control group on the 8th day after surgery before testosterone treatment (p<0.01), and there was no significant difference between experimental group and model control group. After 3 weeks of treatment, the body weight of experimental group and model control group was still significantly lower than sham operation group and blank control group (p<0.01), and there was no significant difference between experimental group and model control group. Three weeks after intramuscular injection of testosterone propionate, serum testosterone concentration in experimental group was significantly higher than three groups (p<0.01, Fig. 1).

**Figure 1.**
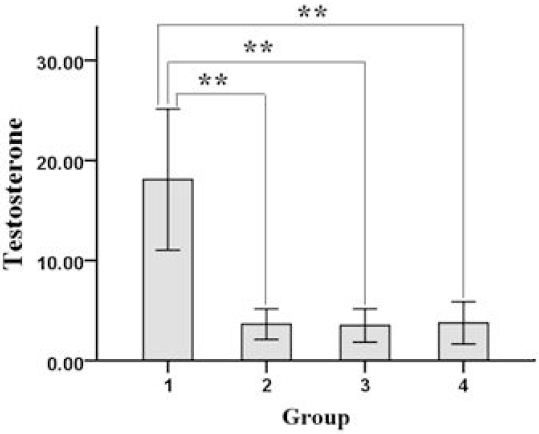
Serum concentration of testosterone in each group of rats. The serum testosterone concentration was determined by radioimmunoassay. 1. experimental group; 2. model control group; 3. sham operation group; 4. blank control group. ** *p* < 0.01.

### Testosterone improved the strength of EDL

The maximal contractile force of EDL was significantly greater in experimental group than in model control group (Fig. 2A). Moreover, the contraction time of EDL was significantly shorter in experimental group than in model control group (Fig. 2B). In addition, the fatigue index of EDL was significantly higher in experimental group than in model control group (Fig. 2C). Collectively, these data indicate that testosterone improved the strength and endurance of EDL.

**Figure 2.**
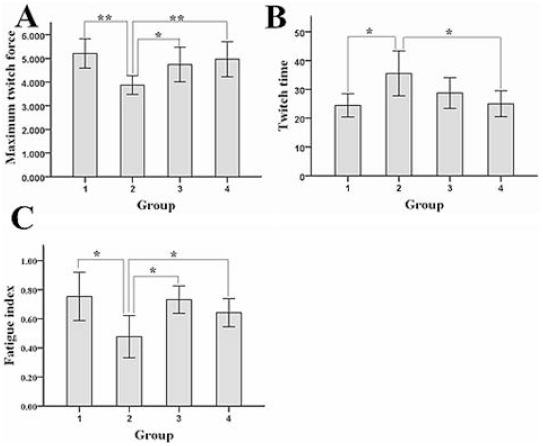
Muscle function of each group of rats. A. Maximum twitch force. B. Twitch time. C. Fatigue index. 1. experimental group; 2. model control group; 3. sham operation group; 4. blank control group. ** *p* < 0.01, * *p* < 0.05.

### Testosterone improved the muscle of EDL

The cross-sectional area of MHC fast muscle fibers of EDL was significantly larger in experimental group than in model control group, while it was also significantly larger in sham operation group and blank control group than in model control group (Fig. 3A). Collectively, these data indicate that testosterone improved the muscle and angiogenesis of EDL.

**Figure 3.**
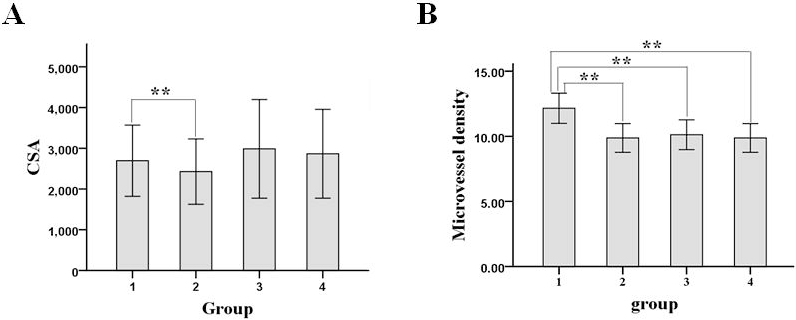
Muscle characteristics of each group of rats. A. Average cross-sectional area (CSA) of fast muscle fiber. 1. experimental group; 2. model control group; 3. sham operation group; 4. blank control group. ** *p* < 0.01.

### Testosterone enhanced the expression of IGF-1 and activated Akt/mTOR signaling in EDL

To reveal the mechanism by which testosterone improved the muscle function of EDL, we detected the expression of IGF-1 in EDL. Western blot analysis showed that IGF-1 protein level in EDL was significantly higher in experimental group than in model control group, sham operation group and blank control group (Fig. 4A). Moreover, the levels of phosphorylated (activated) AKT and mTOR in EDL were significantly higher in experimental group than in control group, sham operation group and blank control group (Fig. 4B). In addition, protein level of fast myosin skeletal heavy chain (MHC) was significantly higher in experimental group than in model control group, and it was also significantly higher in sham operation group and blank control group than in model control group (Fig. 4C).

**Figure 4.**
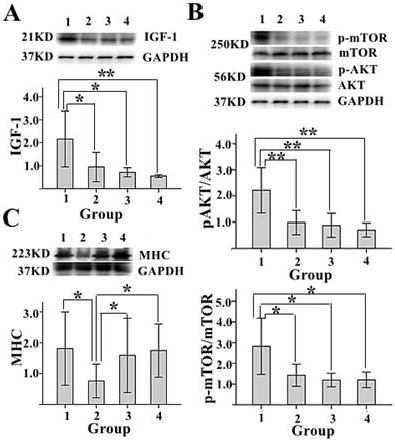
The status of IGF-1/AKT pathway in EDL of each group of rats. A. The level of IGF-1. B. The levels of p-AKT and p-mTOR. C. The level of fast myosin skeletal heavy chain (MHC). GAPDH was loading control. 1. experimental group; 2. model control group; 3. sham operation group; 4. blank control group. ** *p* < 0.01, * *p* < 0.05.

### Testosterone did not cause pathological changes in the testis and prostate

HE staining of the testes in experimental group showed no significant atrophy compared with model control group, sham operation group and blank control group (Fig. 5). HE staining of the prostate in experimental group showed no significant hyperplasia compared with model control group, sham operation group and blank control group (Fig. 6).

**Figure 5.**
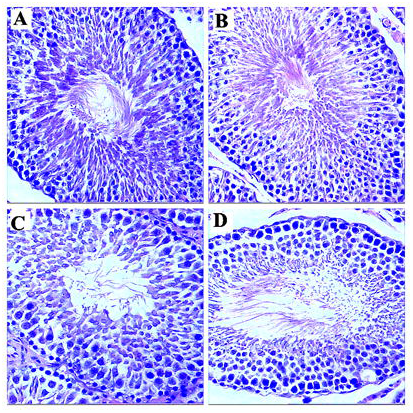
Histological analysis of the testicle in each group of rats. A. experimental group; B. model control group; C. sham operation group; D. blank control group.

**Figure 6.**
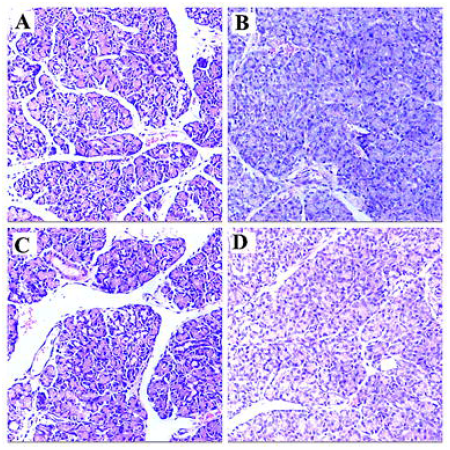
Histological analysis of the prostate in each group of rats. A. experimental group; B. model control group; C. sham operation group; D. blank control group.

## Discussion

Sepsis is one of the major causes of acquired muscle weakness in either human or rats, with a high incidence and severe symptoms of ICUAW. According previous studies, sepsis especially severe sepsis can lead to muscle wasting, with skeletal muscle atrophy, decreased strength and endurance as typical clinical manifestation. ^14,22^ One of the major mechanisms is that sepsis can cause fast degradation of skeletal muscle proteins, for example, skeletal myosin degradation is observed in sepsis models with the characteristic manifestation of ICUAW ^14,22,23^. Therefore, sepsis rat model is a commonly used ICUAW animal models, and the cecal ligation perforation method can not only simulate clinical manifestations of sepsis caused by intestinal perforation and intestinal necrosis but also adjusts the severity of sepsis by controlling the length of the cecal ligation.^16^ In this study, we chose rat sepsis model as a suitable animal model to develop new drug treatment for ICUAW.

Sepsis, especially severe sepsis, can lead to a significant decrease in serum testosterone levels in either patients or experimental animals, which is currently thought to be caused by the suppression of the hypothalamic-pituitary-gonadal axis by systemic inflammatory response syndrome.^17^ In view of the powerful role of testosterone in enhancing exercise strength and endurance, we wondered whether testosterone can improve the symptoms of muscle atrophy following sepsis.

Testosterone propionate is an artificial testosterone preparation that has a long half-life after intramuscular injection. In the past, testosterone propionate has been used to treat severe burns and elderly patients who have conditions of frailty. The typical clinical manifestations of ICUAW include muscle weakness, muscular atrophy and decreased exercise endurance, very similar to sarcopenia caused by other factors. In this study, we found that testosterone propionate promoted maximum contraction force, fatigue index, average cross-sectional area of muscle fiber of rats with sepsis-acquired muscle weakness after short term application. These data indicate that testosterone could enhance the skeletal muscle of rats with sepsis-acquired muscle weakness in strength, endurance, and volume. Moreover, the contraction time of EDL in experimental group was significantly shorter than that in model control group, indicating that skeletal muscle was more excitatory and responsive after testosterone treatment. In addition, rapid myosin degradation is one characteristic of ICUAW, and testosterone significantly increased the expression of rapid myosin in the skeletal muscle of rats with sepsis, which indicates that testosterone could improve the muscle in condition of ICUAW. Unlike some failed intervention previously with growth hormone, this study showed positive effects of testosterone on ICUAW therapy for the first time.

IGF-1/AKT pathway is one of the main signaling pathways that regulate skeletal muscle anabolism and skeletal muscle function.^18^ In this study, serum testosterone level, IGF-1 protein level, phosphorylated AKT and mTOR levels and rapid myosin expression level all significantly increased in EDL of experimental group. These changes suggest that the IGF-1/AKT signaling pathway might be one of the potential mechanisms by which testosterone propionate enhances muscle strength, endurance and muscle volume in septic rats. Furthermore, testosterone propionate has androgen-like effects. Long-term usage of exogenous androgen may inhibit male gonads, and common complications include testicular atrophy and prostatic hyperplasia, which severely limits its clinical application.^19^ Therefore, medicinal plants have been recently exploited to screen new agents with androgen-like effects. ^20,21^ Based on histological characteristics of the prostate and testis of rats in experimental group and other groups, we found no significant pathological changes of the prostate and testis in experiment group, which indicates that testosterone propionate may be safe for short-term use.

However, we should point out some limitations of the intervention with testosterone propionate. For instance, rats in the experimental group were more aggressive compared to control group, such as fighting, baiting, and they had more active sexual behavior. These results might be side effects of testosterone propionate treatment, and the dose and duration of testosterone propionate treatment need further studies.

In summary, our study provides strong evidence that testosterone propionate can significantly improve skeletal muscle strength, endurance and volume in septic rats, and the mechanism may be related to the activation of IGF-1/AKT pathway. Moreover, testosterone propionate with short duration does not cause testicular atrophy and prostate hyperplasia in septic rats. Therefore, our results suggest that testosterone propionate is a potential treatment for muscle malfunction in ICUAW patients.

## Author contribution

Jinlong Wang designed the study, performed some experiments and wrote the manuscript. Tong Wu performed some experiments and approved the manuscript.

## Disclosure

The authors report no conflicts of interest in this work.

## Funding

This study received no any funding.

## References

1. Bolton CF. Neuromuscular manifestations of critical illness. Muscle Nerve. 2005;32(2):140–163.

2. Schefold JC, Bierbrauer J, Weber-Carstens S. Intensive care unit-acquired weakness (ICUAW) and muscle wasting in critically ill patients with severe sepsis and septic shock. J Cachexia Sarcopenia Muscle. 2010;1(2):147–157.

3. Khan J, Harrison TB, Rich MM, Moss M. Early development of critical illness myopathy and neuropathy in patients with severe sepsis. Neurology. 2006;67(8):1421–1425.

4. Latronico N, Shehu I, Seghelini E. Neuromuscular sequelae of critical illness. Curr Opin Crit Care. 2005;11(4):381–390.

5. Jeschke MG, Finnerty CC, Suman OE, Kulp G, Mlcak RP, Herndon DN. The effect of oxandrolone on the endocrinologic, inflammatory, and hypermetabolic responses during the acute phase postburn. Ann Surg. 2007;246(3):351–360.

6. Krause Neto W, Silva WA, Ciena AP, Nucci RAB, Anaruma CA, Gama EF. Effects of Strength Training and Anabolic Steroid in the Peripheral Nerve and Skeletal Muscle Morphology of Aged Rats. Front Aging Neurosci. 2017;9:205.

7. Serra C, Tangherlini F, Rudy S, et al. Testosterone improves the regeneration of old and young mouse skeletal muscle. J Gerontol A Biol Sci Med Sci. 2013;68(1):17–26.

8. Fatani SH, Abdelbasit NA, Al-Amodi HS, Mukhtar MM, Babakr AT. Testosterone, obesity, and waist circumference as determinants of metabolic syndrome in Saudi women. Diabetes Metab Syndr Obes. 2018;11:175–181.

9. Oki K, Law TD, Loucks AB, Clark BC. The effects of testosterone and insulin-like growth factor 1 on motor system form and function. Exp Gerontol. 2015;64:81–86.

10. Sipilä S, Narici M, Kjaer M, et al. Sex hormones and skeletal muscle weakness. Biogerontology 2013;14(3):231–245.

11. Katsuji A. Sex steroid hormones and skeletal muscle. Jpn J Phys Fitness Sports Med. 2016;65(5):455–462.

12. Ghanim H, Dhindsa S, Batra M, et al. Effect of Testosterone on FGF2, MRF4, and 14 Myostatin in Hypogonadotropic Hypogonadism: Relevance to Muscle Growth. J Clin Endocrinol Metab. 2019;104(6):2094–2102.

13. Bhasin S, Storer TW, Berman N, et al. The effects of supraphysiologic doses of testosterone on muscle size and strength in normal men. N Engl J Med. 1996;335(1):1–7.

14. Rossignol B, Gueret G, Pennec JP, et al. Effects of chronic sepsis on contractile properties of fast twitch muscle in an experimental model of critical illness neuromyopathy in the rat. Crit Care Med. 2008;36(6):1855–1863.

15. Rannou F, Pennec JP, Rossignol B, et al. Effects of chronic sepsis on rat motor units: experimental study of critical illness polyneuromyopathy. Exp Neurol. 2007;204(2): 741–747.

16. Williams AB, Decourten-Myers GM, Fischer JE, Luo G, Sun X, Hasselgren PO. Sepsis stimulates release of myofilaments in skeletal muscle by a calcium-dependent mechanism. FASEB J. 1999;13(11):1435–1443.

17. Martín AI, Priego T, López-Calderón A. Hormones and Muscle Atrophy. Adv Exp Med Biol. 2018;1088:207–233.

18. IGF-1 prevents simvastatin-induced myotoxicity in C2C12 myotubes. Bonifacio A, Sanvee GM, Brecht K, et al. Arch Toxicol. 2017;91(5):2223–2234.

19. Kim KS, Yang HY, Chang SC, et al. Potential repositioning of GV1001 as a therapeutic agent for testosterone-induced benign prostatic hyperplasia. Int J Mol Med. 2018;42(4):2260–2268.

20. Villa-Hernández JM, García-Ocón B, Sierra-Palacios EC, Pelayo-Zaldivar C. Molecular biology techniques as new alternatives for medicinal plant identification. Phyton, Int J Exp Botany 2018;87:72–78.

21. Montes FQ, Vázquez-Hernández A, Fenton-Navarro B. Active compounds of medicinal plants, mechanism for antioxidant and beneficial effects. Phyton, Int J Exp Botany 2019;88:1–10.

22. Bird SJ, Rich MM: Critical illness myopathy and polyneuropathy. Curr Neurol Neurosci Rep 2002; 2:527–533.

23. Williams AB, Decourten-Myers GM, Fischer JE, et al: Sepsis stimulates release of myofilaments in skeletal muscle by a calciumdependent mechanism. Faseb J 1999; 13:1435–1443.

